# Intraspecific variation shapes community-level behavioural responses to urbanisation in spiders: from traits to function

**DOI:** 10.1101/076497

**Authors:** Maxime Dahirel, Jasper Dierick, Maarten De Cock, Bonte Dries

## Abstract

1. Approaches based on functional traits have proven especially valuable to understand how communities respond to environmental gradients. Until recently, they have, however, often ignored the potential consequences of intraspecific trait variation (ITV). This position becomes potentially more problematic when studying animals and behavioural traits, as behaviours can be altered very flexibly at the individual level to track environmental changes.
2. Urban areas are an extreme example of human-changed environments, exposing organisms to multiple, strong, yet relatively standardized, selection pressures. Adaptive behavioural responses are thought to play a major role in animals’ success or failure in these new environments. The consequences of such behavioural changes for ecosystem processes remain understudied.
3. Using 62 sites of varying urbanisation level, we investigated how species turnover and ITV influenced community-level behavioural responses to urbanisation, using orb web spiders and their webs as models of foraging behaviour.
4. ITV explained around 30% of the total trait variation observed among communities. Spiders altered their web-building behaviour in cities in ways that increase the capture efficiency of webs. These traits shifts were partly mediated by species turnover, but ITV increased their magnitude. The importance of ITV varied depending on traits and on the spatial scale at which urbanisation was considered. Available prey biomass decreased with urbanisation; the corresponding decrease in prey interception by spiders was less important when ITV in web traits was accounted for.
5. By facilitating trait-environment matching despite urbanisation, ITV thus helps communities to buffer the effects of environmental changes on ecosystem functioning. Despite being often neglected from community-level analyses, our results highlight the importance of accounting for intraspecific trait variation to fully understand trait responses to (human-induced) environmental changes and their impact on ecosystem functioning.

## Introduction

Trait-based approaches to community ecology have provided a valuable framework to understand how communities respond to environmental gradients, and the consequences for ecosystem functioning (Lavorel & Garnier 2002; Moretti *et al.* 2009; Cornwell & Ackerly 2009; Lavorel *et al.* 2011; Dray *et al.* 2014; Jung *et al.* 2014; Simons, Weisser & Gossner 2016). Community-level trait responses have generally been investigated by focusing on species turnover, ignoring the potential effects of intraspecific trait variation (ITV) due to plasticity and/or evolutionary change. This has been done both for practical reasons and under the commonly-held assumption that ITV is negligible relative to between-species variation (Albert *et al.* 2011; Violle *et al.* 2012). An increasing number of studies shows, however, that ITV may represent a non-negligible part of the total trait variation observed within and among communities (reviewed in Siefert *et al.* 2015), and that correctly accounting for it may greatly change the strength of estimated community-level trait shifts along environmental gradients (Lepš *et al.* 2011; Jung *et al.* 2014).

Compared to morphological traits typically used in trait-based (plant) community ecology, animal behaviours are usually seen as more flexible (Pigliucci 2001; Duckworth 2008; Sih *et al.* 2010), which would allow for a greater importance of ITV to community-level responses. However, and despite the fact that intraspecific variation in behaviour can have wide-ranging impacts on community dynamics and ecosystem functioning (Modlmeier *et al.* 2015), the relative effects of inter-*versus* intraspecific behavioural variation are rarely compared. We expect that intraspecific variation in behaviour will be even more relevant for communities experiencing rapid and strong environmental changes, such as human-induced rapid environmental changes (HIREC *sensu* Sih *et al.* 2010), as behavioural flexibility is the first and fastest line of response of animals in these contexts (Sih *et al.* 2010; Wong & Candolin 2015). Both adaptive and maladaptive changes in behaviours in response to HIREC have been recorded within numerous taxa, and many questions remain on the determinants of these changes, and their impact on community-level processes and ecosystem functioning (Wong & Candolin 2015).

Urban areas now concentrate more than 50% of the world population on less than 3% of the world land surfaces, with these two numbers predicted to increase in the near future (Seto, Güneralp & Hutyra 2012; Liu *et al.* 2014; United Nations Population Division 2015). The strong and multiple environmental changes associated with urbanisation all contribute to create evolutionary novel environments in which many species are unable to fit and disappear, yet some manage to exploit these new opportunities and persist or even proliferate (McKinney 2006, 2008; Croci, Butet & Clergeau 2008; Aronson *et al.* 2014; Knop 2016). Species able to pass such a strong and multivariate external filter (*sensu* Violle *et al.* 2012) are expected to possess consistently different functional trait values than those that cannot and are excluded (e.g. Croci *et al.* 2008), which should lead to shifts in mean community-level trait values in urbanized environments.

Orb-web weaving spiders (Arachnida; Araneae; main families: Araneidae and Tetragnathidae) are ubiquitous generalist predators present in many natural and human-altered terrestrial ecosystems (Roberts 1993; Sattler *et al.* 2010; Foelix 2010). Orb-web design and size are highly variable both among and within species (Bonte *et al.* 2008; Sensenig, Agnarsson & Blackledge 2010). Differences between species, populations and individuals in e.g. silk investment, web positioning, capture area, or mesh width have been linked to prey availability, and are thought to reflect adaptive decisions aiming to maximize the benefits/costs ratio of trap-building in different contexts (Sherman 1994; Blackledge & Eliason 2007; Bonte *et al.* 2008; Blamires 2010; Scharf, Lubin & Ovadia 2011; Eberhard 2013). Orb webs can therefore be seen as high-resolution and easy to access archives of foraging decisions (Sherman 1994), greatly facilitating the acquisition of *in situ* behavioural data for all species in a community. The prey control function of spider communities often goes beyond mere consumption, as webs may capture more prey than are actually eaten (Riechert & Maupin 1998), and the mere presence of silk can deter herbivorous insects from feeding on a given plant (Hlivko & Rypstra 2003; Rypstra & Buddle 2013). Juvenile orb web spiders can disperse over large distances by ballooning, yet their movement is generally much more limited after settlement (Lubin, Ellner & Kotzman 1993; Foelix 2010); spiders are therefore expected to be strongly affected by local conditions (Sattler *et al.* 2010; but see Lowe, Wilder & Hochuli 2014).

Here we used orb-web spiders and urban ecosystems to test the hypothesis that ITV has a significant role in shaping community-level behavioural responses to environmental changes and their consequences for ecosystem functioning, and that this impact varies with the spatial scale of environmental change. We predicted (1) that urbanisation would negatively influence prey biomass availability, (2) that the average spider web-building strategy would shift to track these prey changes, with a predominant contribution of ITV to observed responses; (3) that the contribution of ITV to overall trait variation would be non-negligible, but less important in body size and traits strongly constrained by body size (e.g. web investment) than in “strict” behavioural traits (e.g. web shape and positioning). Spider body size was also expected to vary due to both changes in prey availability and higher temperature in cities (the heat island effect; Pickett *et al.* 2001), the latter favouring larger spiders (Entling *et al.* 2010), with potential constraining effects on the range of available web-building strategies (Gregorič, Kuntner & Blackledge 2015). Finally, we investigated the effect of intraspecific variation in web-building on changes in a measure of ecosystem functioning, namely the prey control potential of orb-web spider communities, across the urbanisation gradient.

## Material and Methods

### Study sites and sampling design

We sampled 62 orb-weaving spider communities across an urbanization gradient in Flanders (Belgium) (Supplementary Figure 1). The vast majority of the Belgian population is concentrated in cities (97.8%; United Nations Population Division 2015). In order to study the effect of urbanization at different spatial scales, site selection was carried out following a stratified 2-step design, using geographic information system (GIS). First, 21 non-overlapping plots (3 × 3 km, hereafter “landscape scale”) were selected to represent three urbanization levels (7 plots by level): “low-urbanisation” plots had less than 3% of their surfaces occupied by buildings and more than 20% by ecologically valuable areas according to the Flanders-wide “Biological Valuation Map” (Vriens *et al.* 2011); “high-urbanisation” plots were defined by a percentage > 10% of built-up surfaces and intermediate areas between 5 and 10%. In a second step, we selected within each plot 3 subplots (200 × 200 m, hereafter the “local scale”), one per urbanization level, this time based on built-up area only. Vegetated areas in selected subplots were grassland-dominated and unforested, with shrubs and low trees (in e.g. gardens, parks or hedgerows). One of these 63 subplots was not sampled in time due to bad weather. Sites belonging to different urbanization levels based on these criteria also differed significantly in estimated population density (higher in high-urbanisation sites at both spatial scales; Supplementary Figure 2) and average temperature (with significantly higher temperatures in urbanised sites; Kaiser, Merckx & Van Dyck 2016).

### Collection of spiders and species determination

Field work took place between August 27 and October 5, 2014. One plot was sampled per day; sampling was organized so there was no significant link between plot-level urbanization and sampling date (ANOVA; *N* = 21 plots, *F*_*2,18*_ = 0.009, *p* = 0.991). Each subplot was explored until no new orb web could be found, and every encountered spider was caught and stored in 70% ethanol. The 2456 adult individuals belonging to 18 species were captured and their cephalothorax width measured as a proxy for body size (species determination under binocular microscope, based on Roberts 1993) (Supplementary Table 1).

### Web characteristics

As collecting web information for all sampled spiders was too time-consuming, 944 webs were randomly analysed across all plots. On average, 5.82 webs (*SD* = 15.86, range: 0 - 72) were sampled per species per combination of local and landscape urbanisation levels (hereafter termed the “urban context”). At the level of the local × landscape urban context, the number of webs analysed per species was proportional to the number of spiders captured (162 possible species - urban context combinations; *N*_*webs*_ = −1.80 + 0.50 × *N*_*spiders*_, *R²* = 0.85).

For each selected web, the following web design parameters were measured in the field: the vertical and horizontal web diameters (divided between the central area, i.e. the free zone around the hub without sticky spirals, and the peripheral capture area), the number of sticky silk spirals (*sensu* Zschokke 1999) along the horizontal and vertical axes, and web height (distance from ground to web centre). From these, we calculated the total length of sticky silk spirals (capture thread length, or CTL) following Venner *et al.* (2001)’s formula, the average mesh size in the capture area, as well as the web capture area surface (total web surface minus central area surface), by considering orb webs as ellipses (Herberstein & Tso 2000).

### Prey availability

Prey characteristics as a function of landscape and local urbanization levels were assessed only in 9 plots located in the region of the city of Ghent using sticky paper traps. In each subplot three traps (analysed area: 100 cm² per trap; Pherobank, Wijk bij Duurstede, Netherlands) were placed at about one meter high (in the present study, the overall mean web height was 89.6 cm, with *SD* = 39.45 cm). Traps were operational during the spider sampling period, and were collected on average after 17.8 days (*SD* = 4.41, range: 8 to 26 days). Because some traps went missing or were destroyed before collection, data were only available for 25 out of the 27 sampled subplots. Preys were counted in the lab and their body length measured to the mm. Prey biomass was approximated using the following equation: dry mass (mg) = 0.04 × body length (mm)^2.26^ (Sabo, Bastow & Power 2002); this applies well to Diptera that are the main prey for orb web spiders in the studied landscapes (Ludy 2007).

### Statistical analysis

All analyses were carried out using R, version 3.2 (R Core Team 2016).

For each spider trait (body size, web height, CTL, capture area surface and mesh size), we investigated shifts in mean values across the urbanization gradient with community-weighted mean values: 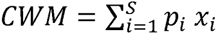, where *S* is the number of species in the community, *x*_*i*_ the relevant trait value of species *i* and *p*_*i*_ its relative abundance in the sample. We used the definitions and methods of Lepš *et al.* (2011) to distinguish the contributions of inter- and intra-specific variation to changes in these mean trait responses across environmental gradients. For each trait at each site we calculated three *CWM* values. “Fixed” averages were based on species mean trait values calculated by averaging over all samples; changes in fixed averages can therefore only reflect the effect of species turnover (changes in species occurrence or relative abundance). “Specific” averages were based on separate species mean trait values for each urban context and thus reflect both species turnover and intraspecific variation in response to the urbanization gradient. The intraspecific trait variation (ITV) effect was defined as the difference between specific and fixed values. In the small minority of species × urban context combinations where no web was sampled at the urban context level (*N* = 68 individuals out of 2456 = 2.77 % of all sampled spiders, range per site: 0 – 16.67% of individuals), we used the same mean trait value for the fixed and specific *CWM* calculations.

Linear models were then fitted to specific, fixed, and ITV values of all traits, allowing us to test for the effect of environment on overall community response and its separate components. We included effects of local- and landscape-scale urbanization level, with an effect of plot identity to account for spatial/temporal clustering. In models for web traits, because most web traits are at least partly correlated with body size (Heiling & Herberstein 1998; Sensenig *et al.* 2010; Gregorič *et al.* 2015), we added the fixed and intraspecific components of body size *CWM* as covariates. Given the “specific” response of a trait results by definition from the addition of the fixed and intraspecific responses, and taking the total variability in “specific” values to be 100%, we used ANOVAs and sum of squares (*SS*) decomposition to partition trait variation into its fixed and intraspecific components (each extracted from their respective model; Lepš *et al.* 2011). This uses the fact that the variance of the sum of two independent variables is equal to the sum of their variances: *SS*_*specific*_ = *SS*_*fixed*_ + *SS*_*ITV*_ (with degrees of freedom held constant across all three models). If fixed and intraspecific components are not independent but covary, the total *SS*_*specific*_ will be higher or lower than expected based on *SS*_*fixed*_ and *SS*_*ITV*_, depending respectively on whether the covariation is positive or negative. We can therefore determine the proportion of total variation due to covariation between inter- and intraspecific responses: *SS*_*cov*_ = *SS*_*specific*_ –*SS*_*fixed*_ – *SS*_*ITV*_. This variation partitioning was done for both the overall variation and the part of the variation explained by each variable introduced in linear models (Lepš *et al.* 2011). Given the high number of tests involving community weighted mean traits, all *p*-values from trait ANOVAs were adjusted using Benjamini & Hochberg (1995)’s method, in order to minimize the false discovery rate. Tukey’s Honest Significant Differences tests were carried out when a significant urbanization effect was found in ANOVAs to identify and quantify differences between urbanization levels.

In addition, we used data on average web characteristics, spider abundance and prey characteristics to quantify the effect of urbanisation on overall prey interception potential of communities, with or without ITV being taken into account. We evaluated the effect of urbanization level on overall subplot-level spider abundance, mean prey size, mean number of prey and *B*, the mean prey biomass caught per day per cm² of sticky trap using generalized linear mixed models with a Poisson family for the former (*N* = 62) and linear mixed models for the latter three (*N* = 25), including a random effect of plot identity to account for site clustering (using the R package lme4, Bates *et al.* 2015). Then, for the 25 communities in which prey were sampled, we evaluated the proportion *P* of the prey biomass *B* expected to actually contact the web and be intercepted: for each web and prey item caught in sticky traps, we assumed the latter would touch the former with a probability = 1 if prey body length > mesh size, with a probability = body length / mesh size otherwise (Evans 2013). The overall daily prey interception potential of a spider community was thus equal to *B* × *P* × spider abundance × web capture surface *CWM*. As above, we calculated specific, fixed and intra-specific values of prey interception potential, and analysed the effects of local and landscape-level urbanisation, controlling for mean spider size and plot identity, on these different components following Lepš *et al.* (2011).

## Results

### Effect of urbanization on spider abundance

The total number of spider per sample was significantly affected by urbanization at the local scale (Wald tests, *χ*^*2*^ = 20.49, df = 2, *p* = 3.55 × 10^−5^) but not at the landscape scale (*χ*^*2*^ = 1.10, df = 2, *p* = 0.58). Spiders were less abundant in communities experiencing locally high and intermediate levels of urbanization, compared to low-urbanization communities (Tukey tests, *p* < 7.08 × 10^−4^ for significant differences; means ± SD = 36.8 ± 8.1, 37.4 ± 6.3 and 44.7 ± 9.6 spiders per 4 ha site, respectively; overall average: 39.6 ± 8.8).

### Trait variation partitioning (Fig. 1, Table 1)

Species turnover alone explained 30.93 (web height) to 78.18% (body size) of the total variance in specific mean values, depending on traits (average: 59.32%), with intraspecific trait variation and covariation between inter- and intra-specific responses making up for the difference. Plot identity accounted for a great part of the variation in body size (44.54%). The turnover component of body size differences among communities explained a substantial part of the overall variation in average mesh size (30.15 %), CTL (35.96 %) and web surface (57.95 %), but not in average web height (0.98 %). The proportion of web trait variation explained by intraspecific variation in body size was much lower (max. 3.0 % for mesh width, average for the four web traits: 1.22 %). Urbanisation (landscape and local scales combined) accounted for between 10.31 (body size) and 75.74 % (web height) of total variation, depending on traits (average: 34.94 %). Based on Lajoie and Vellend (2015), we calculated the relative contribution of ITV to responses to urbanisation: SS_ITV(urbanisation)_ / (SS_ITV (urbanisation)_+ SS_fixed (urbanisation)_), excluding the covariation between the two components SS_cov (urbanisation)_. It ranged from 44.69% (mesh width) to 90.31% (web surface) (average across the five tested traits: 67.26 %).

**Figure 1.**
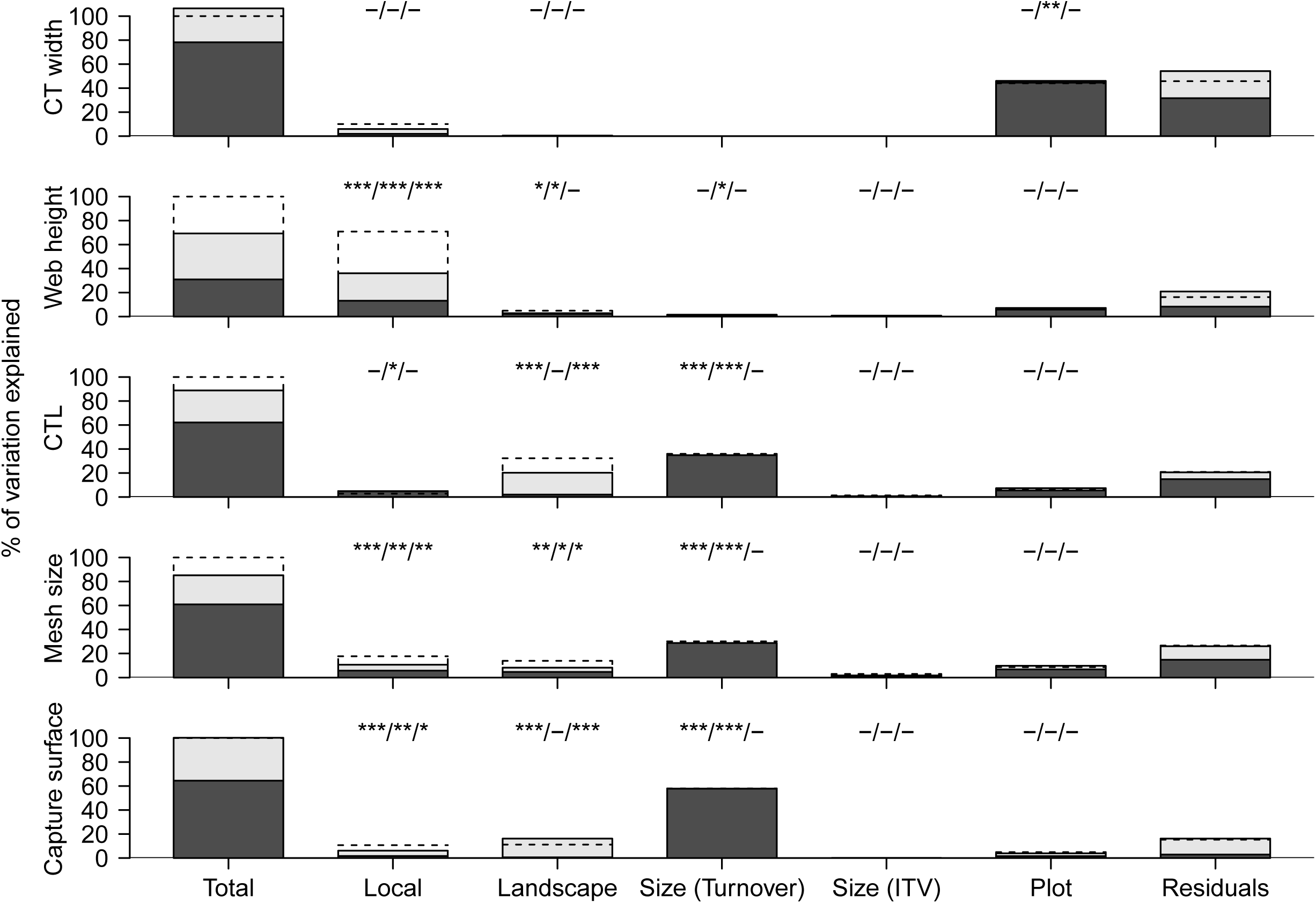
Partitioning of the variation in community-weighted mean trait values, following Lepš *et al.* (2011). Dark grey parts of each bar correspond to species turnover effects, light grey to intraspecific variability (ITV) effects. Dotted lines denote total variation (in “specific” values, i.e. including both turnover and ITV). Differences between the dotted lines and the sum of turnover and ITV effects correspond to the effect of covariation. Asterisks denote significant effects on the habitat-specific/turnover/ITV components, in that order (ANOVAs; *: *p* < 0.05, **: *p* < 0.01, ***: *p* < 0.001 after false discovery rate adjustments; full details of statistical tests are presented in Table 1).

**Table 1.**
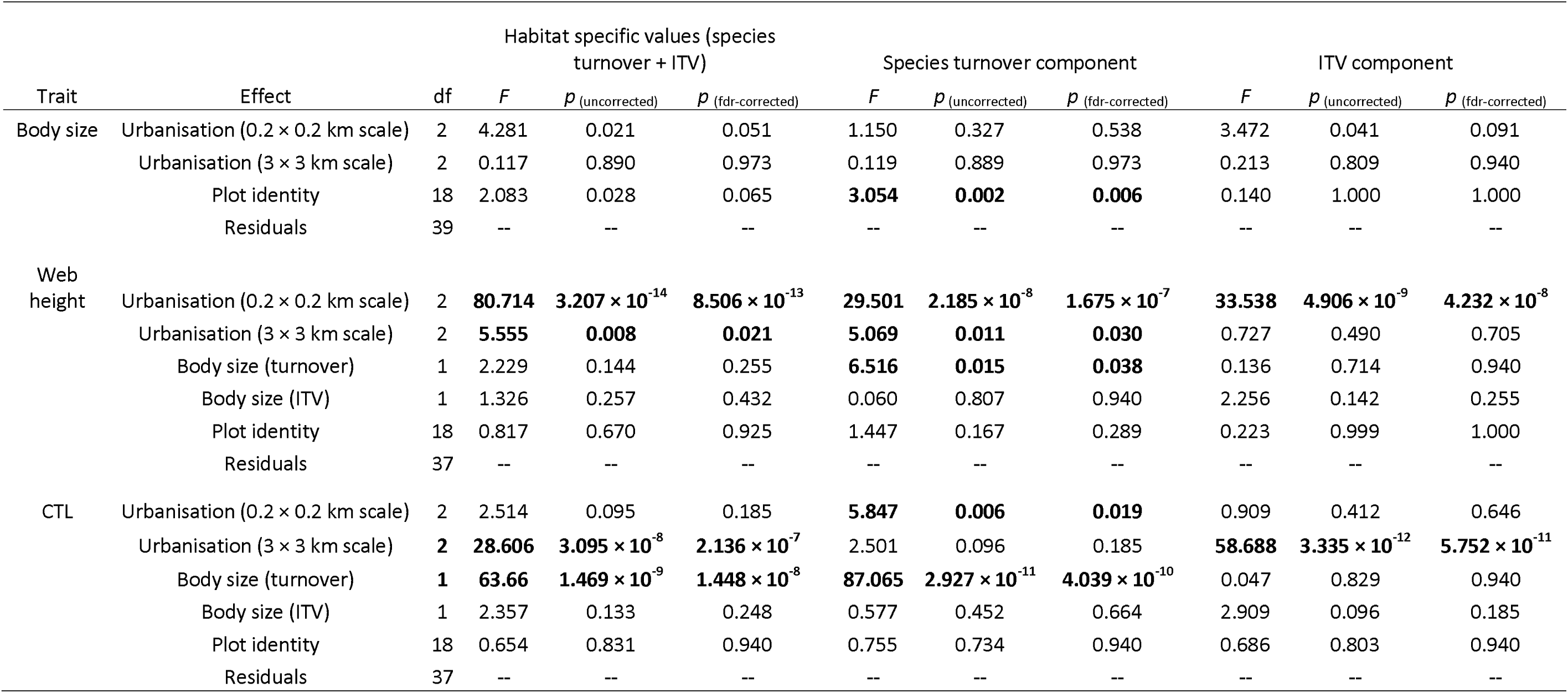
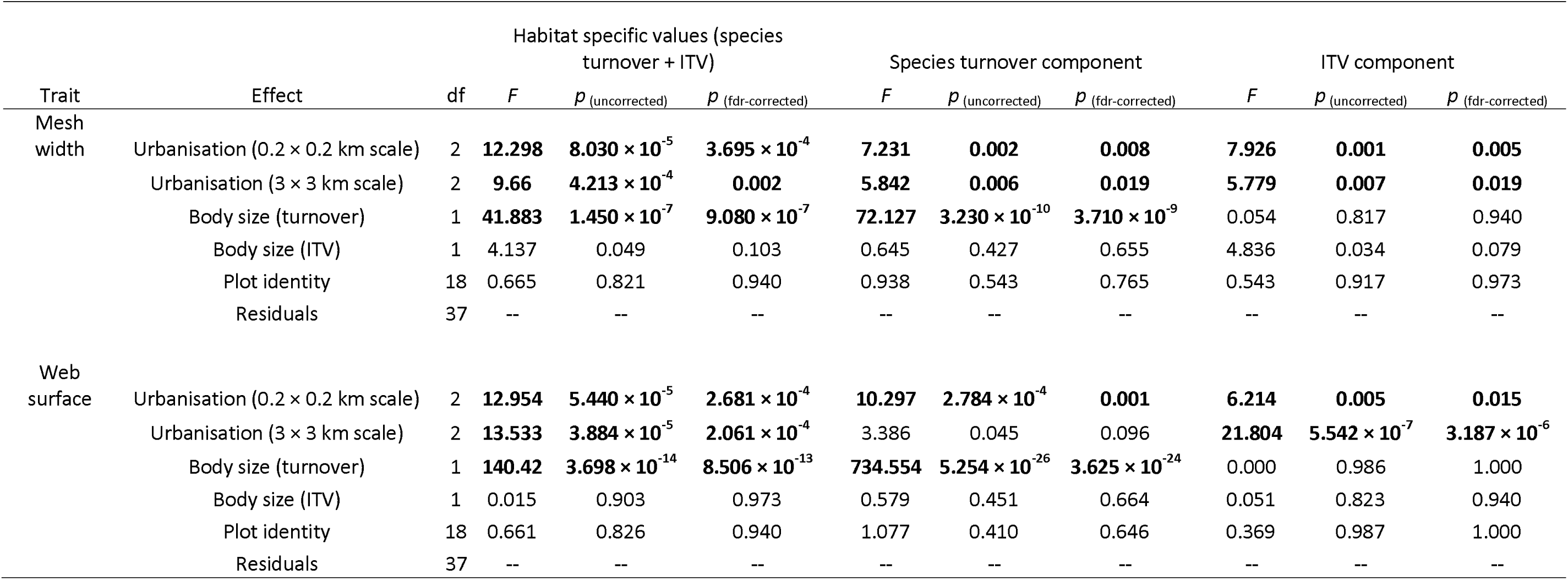
Results from ANOVAs linking variation in spider community-weighted mean traits to urbanisation. The erall trait responses (habitat-specific values), the turnover component and the intraspecific variation component are analyzed separately (Lepš *et al.* 2011). Significant effects (*p* < 0.05 after correcting for false discovery rate following Benjamini & Hochberg 1995) are in bold.

### Effect of urbanization on community-weighted mean trait values (Fig. 2, Table 1)

No significant effect of local or landscape-level urbanisation was detected on either specific mean body size or its separate turnover and intraspecific variation components after *p*-value correction.

**Figure 2.**
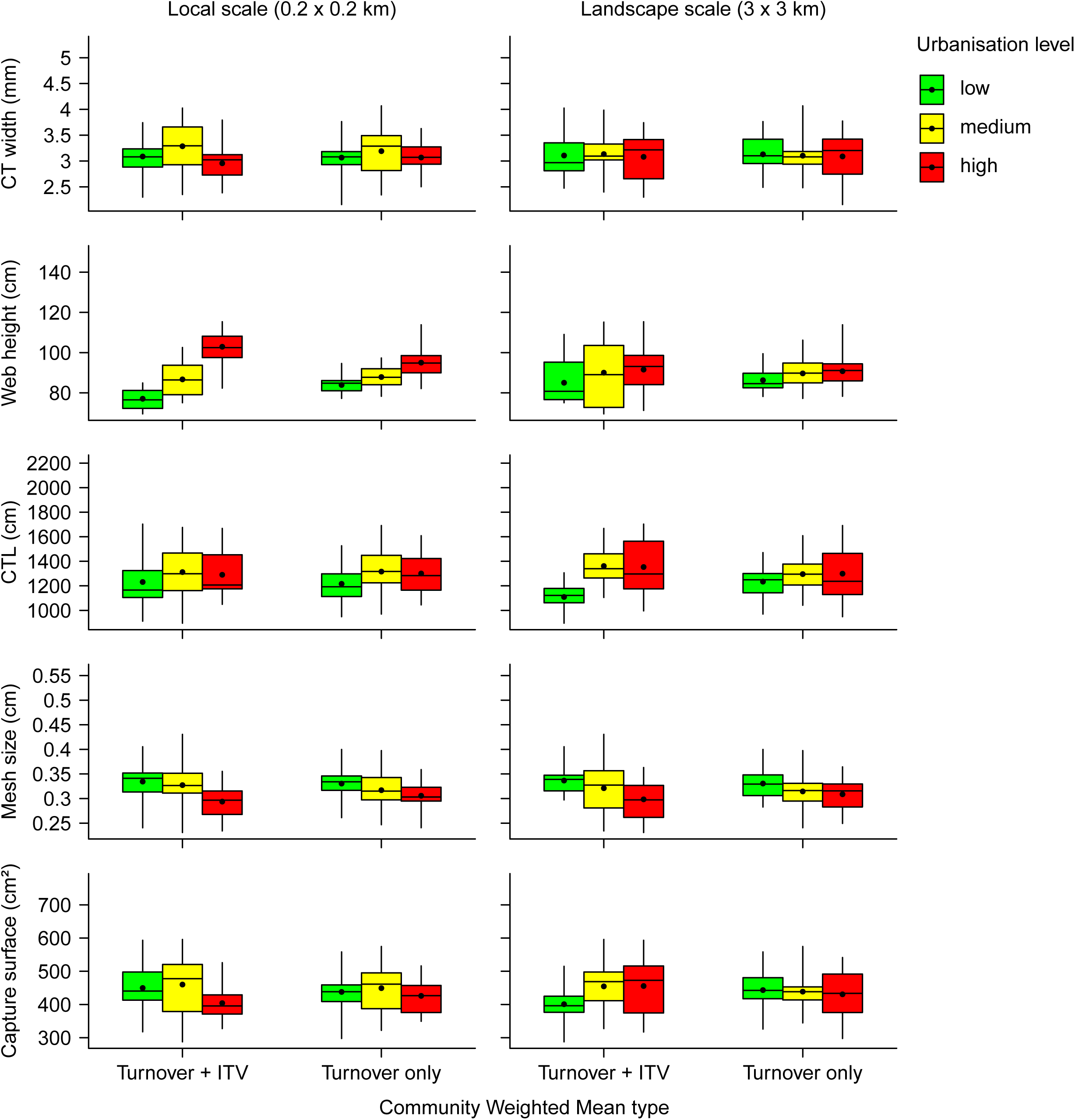
Community weighted mean values of the five spider traits under study, as a function of the local (left column) or landscape (right column) level of urbanization. Boxes indicate the first and third quartiles, whiskers the range. Horizontal lines inside boxes indicate medians, dots denote means. Relevant statistical tests are presented in Table 1.

Spiders built their webs significantly higher as the local level of urbanisation increased, whether one considered specific values or their separated turnover and ITV components (Tukey HSD tests, *p*-values for all pairwise comparisons < 0.03). At the landscape scale, webs were also built significantly higher in highly urbanised communities when compared to more natural sites; this effect was observed with specific values (Tukey HSD tests, *p* = 0.008) and turnover components (*p* = 0.011), but not with ITV values.

Regarding web investment, specific and ITV-only, but not turnover-only, values of CTL were, at the landscape scale, significantly higher in highly and moderately urbanised sites when compared to low-urbanisation communities (Tukey HSD tests, *p* < 4.38 × 10^−7^). At the local scale, the same differences were found, but only on the turnover component (Tukey HSD tests, *p* = 0.027).

All measures of mesh width variation (specific, turnover-only and ITV-only) were significantly affected by urbanisation at both spatial scales. In all cases, mesh width values were significantly lower in highly urbanised sites when compared to low-urbanisation communities (Tukey HSD tests, *p* < 0.021), moderately urbanised sites exhibiting intermediate values.

Mean web surface values were also significantly affected by urbanisation but, contrary to other web traits, here the effects of local and landscape-level urbanisation went in opposite directions in some cases. At the local 200 × 200 m scale, specific and ITV-only values of web surface were significantly lower in highly urbanised sites when compared to low-urbanisation communities (Tukey HSD tests, *p* < 0.010), moderately urbanised sites exhibiting intermediate values. For the turnover-only component, the only significant difference was between highly and moderately urbanised sites, the former having here again the lowest values (Tukey HSD tests, *p* = 1.70× 10^−4^, the *p*-value for the high-/low-urbanisation pairwise comparison being equal to 0.061). At the landscape scale on the other hand, specific and ITV values of web surface were significantly higher in highly urbanised sites when compared to low-urbanisation communities (Tukey HSD tests, *p* < 1.55 × 10^−4^). The turnover component, on the other hand, presented significantly lower values in highly urbanised sites (Tukey HSD test, p = 0.037).

### Effect of urbanisation on prey characteristics and availability

The number of prey caught daily per cm² of trap did not differ significantly between urbanisation levels (Wald tests, p > 0.05; overall mean ± SD: 0.06 ± 0.02). The body length of the average prey and the estimated biomass caught daily per cm² in traps were significantly different between urbanisation levels both at the local (Wald tests, *χ^2^* = 9.62, df = 2, *p* = 8.15 × 10^−3^ and *χ*^*2*^ = 12.00, df = 2, *p* = 2.48 × 10^−3^, respectively) and landscape scale (*χ*^*2*^ = 8.17, df = 2, *p* = 0.02 and *χ*^*2*^ = 8.07, df = 2, *p* = 0.02, respectively). For both variables and both spatial scales, highly urbanised sites presented lower values than natural habitats (Fig. 3).

**Figure 3.**
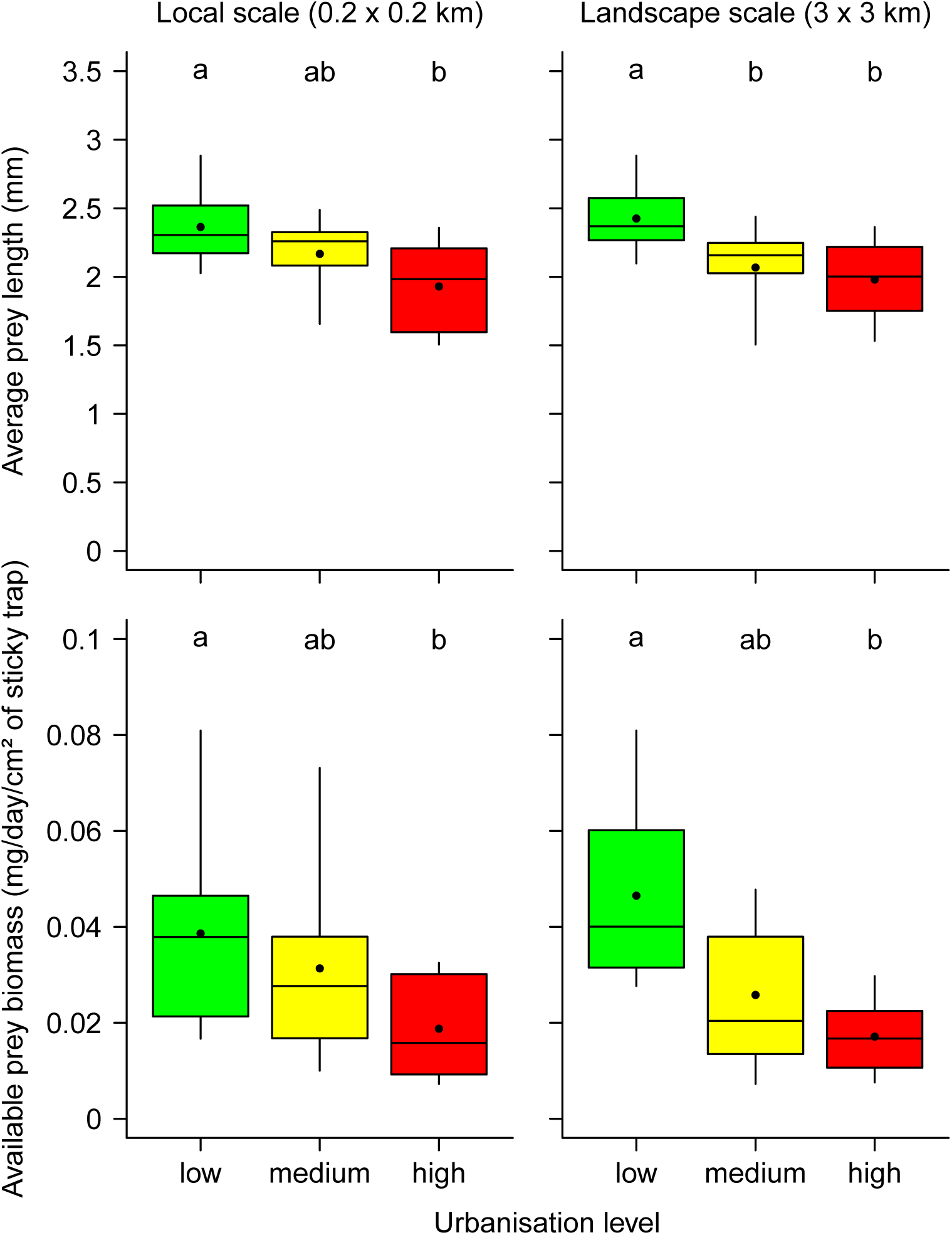
Characteristics of prey caught in sticky traps (top: average body length; bottom: biomass.day^−1^.cm^−2^) as a function of local (left) and landscape-scale (right) level of urbanization. Boxes indicate the first and third quartiles, whiskers the range. Horizontal lines inside boxes indicate medians, dots denote means. Within a subplot, different letters denote significant differences between urbanization levels (p < 0.05, Tukey tests) based on the fixed effects of linear mixed models.

### Effect of urbanization on the prey interception potential of spider communities

The ITV component explained only a small part (*SS*_*ITV*_ =7.0%) of the between-community variance in estimated daily intercepted prey biomass, but there was a substantial negative covariation with the species turnover component (*SS*_*cov*_ = −31.9%, Pearson’s *r* = −0.54, *p* = 0.005) (Fig. 4, top). For both specific (turnover + ITV) and fixed (turnover-only) estimates, and at both spatial scales, there was a significant decline in prey interception potential with increased urbanisation (Fig. 4, bottom; ANOVAs, Supplementary Table 2; Tukey HSD tests, p < 0.01 for comparisons between high and low-urbanisation sites). However, there was a significant increase in ITV values with increased landscape-level urbanisation (ANOVA, *F*_*2,12*_= 20.61, *p* = 1.31 × 10^−4^; Fig. 5). This effect means the decrease in community prey interception when going from low to high urbanisation landscapes is on average 110.12 mg/day (95% CI: 52.56 - 167.68) less important when taking ITV into account, compared to estimates ignoring it (Tukey HSD test on the ITV component, *p* = 7.02 × 10^−4^, Fig. 5). There was no significant effect of the local level of urbanisation on the ITV component of prey interception (ANOVA, *F*_*2,12*_ = 0.06, *p* = 0.94).

**Figure 4.**
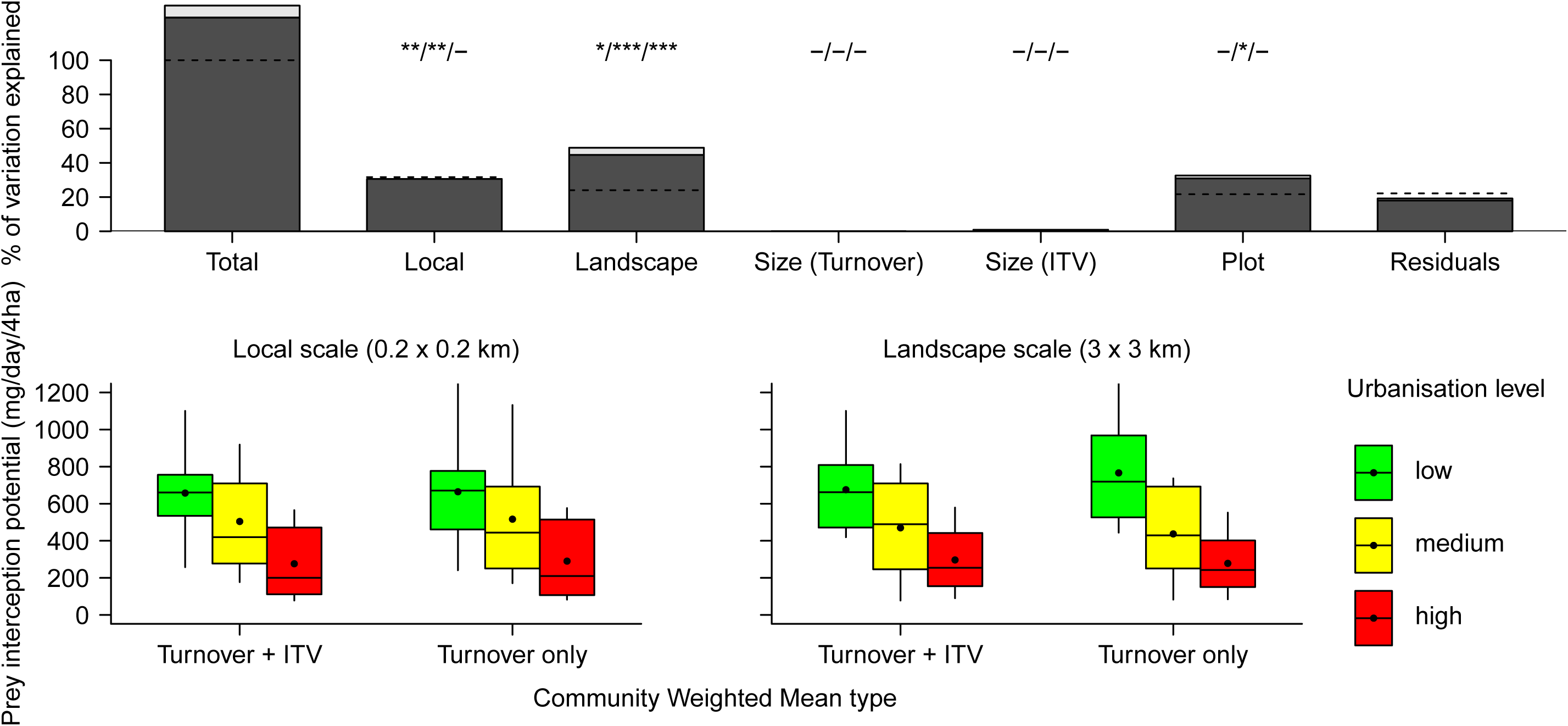
(top) Partitioning of the variation in daily prey interception of spider communities. See Figure 1 for legend details. (bottom) Prey interception potential as a function of local (left) or landscape (right) level of urbanization. Boxes indicate the first and third quartiles, whiskers the range. Horizontal lines inside boxes indicate medians, dots denote means. Relevant statistical tests are presented in Supplementary Table 2.

**Figure 5.**
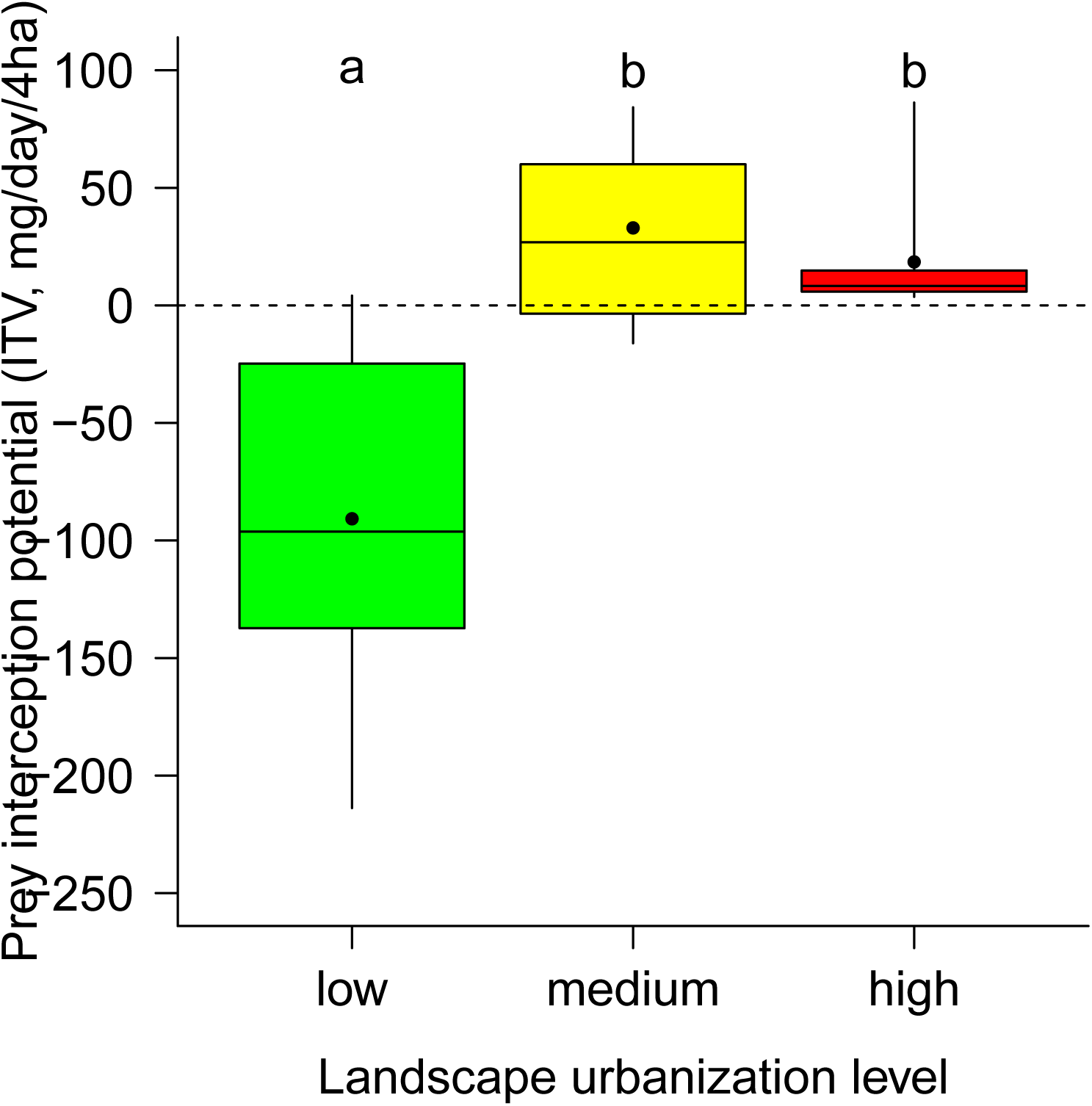
Effect of landscape-scale urbanisation on the intraspecific component of daily prey interception potential. See Fig. 2 for legend details. Levels with different letters are significantly different (p < 0.05, Tukey tests)

## Discussion

Intraspecific trait variation (ITV) contributed substantially to observed between-communities variation in mean trait values (on average around a third, in line with values found in plants; Siefert *et al.* 2015). Urbanisation had negative impacts on both prey biomass availability and spider abundance. ITV generally increased the magnitude of trait changes in response to urbanisation, though the strength of this effect depended on the trait and spatial scale considered. By contrast, it dampened changes in a metric of ecosystem functioning, the potential biomass of prey caught daily.

Based on the common assumption that behaviours are more flexible than morphological traits (Pigliucci 2001; Duckworth 2008), we hypothesized that ITV would be quantitatively more important in “purely” behavioural traits than in web traits more constrained by body size, or body size itself. This is only partly validated by our results. On the one hand, web height, the behavioural trait that was the least constrained by body size (Fig. 1), had the strongest contribution of ITV to among-communities differences, compared to other web traits or body size (Fig. 1). However, the relative contribution of ITV to changes along the urbanisation gradients is more important for CTL and capture surface, which are more constrained by body size (Fig. 1). The fact that the two main components of foraging strategy, namely web-building behaviour and web energetic investment (*sensu* Sherman 1994), appear to be flexible in orb-web spiders means that intraspecific variation should be an important element to understand how spider traits respond to urbanisation.

Responses to urbanisation varied among traits (Fig. 2). No clear effect of urbanisation on community-averaged body size was detected. Population- and community-level changes in orb web spider body size have been observed in response to several of the individual environmental changes associated with urbanisation (Miyashita, Shinkai & Chida 1998; Mayntz, Toft & Vollrath 2003; Opell, Berger & Shaffer 2007; Entling *et al.* 2010), and body size changes with urbanisation have been reported in zooplankton communities, using the same plots as the present study (Kristien Brans et al., unpublished data). While the decrease in prey biomass availability (Fig. 3) may limit growth (Mayntz *et al.* 2003), the heat island effect is expected to favour larger spiders, due to higher temperatures leading to increased metabolism and longer periods available for development (Entling *et al.* 2010; Lowe *et al.* 2014). The absence of overall shift in body size may therefore result from opposite effects of different environmental gradients (possibly on different species) compensating each other, rather than from the absence of actual response.

By contrast, all studied web traits responded to urbanisation. Spiders built on average their webs higher as the urbanisation level increased. This effect was present at both spatial scales, and driven by both species turnover and ITV (although only at the local scale for the latter). Although the precise drivers are not well understood, web height is often dependent on surrounding habitat characteristics such as prey flying height or cover from predators (Herberstein 1997; Blamires, Thompson & Hochuli 2007; Foelix 2010). The presence of high human-built structures, e.g. street lights, might also provide new anchoring points and favour spiders building their webs higher in cities, both at inter- and intra-specific levels.

Available prey biomass decreased as urbanisation increased at both spatial scales, due to a decrease in prey body size, but not prey abundance (Fig. 3). This may be driven by the loss of potential habitat and food resources due to habitat degradation, loss or fragmentation during urbanisation (Smith *et al.*; Oliveira *et al.* 2016; Renauld *et al.* 2016). Contrary to e.g. vertebrates (Fischer *et al.* 2012), orb web spiders cannot switch partly or completely to anthropogenic food sources; urban spider communities thus face strong pressures to build webs adapted to the new prey spectrum. Spiders built webs with smaller mesh widths as urbanisation increased (Fig. 2). This effect was present and quantitatively similar at both spatial scales (Fig. 2) and was both driven by species turnover and intraspecific variability. Mesh width is highly variable between species (Sensenig *et al.* 2010), but also very plastic; individual spiders are able to change this parameter from one web to the next, based on current prey availability and characteristics (Schneider & Vollrath 1998; Blamires 2010). Webs with smaller mesh widths may help maximize prey capture in urban contexts both by facilitating the interception of smaller insects that would pass through looser webs (Sandoval 1994), and/or by increasing the number of contact points between webs and prey, particularly large preys, making them less likely to escape (Blackledge & Zevenbergen 2006). Assuming spiders only have a finite amount of silk to allocate to webs, a small mesh width must be traded off against a smaller web capture surface (Eberhard 2013). Larger webs can intercept more prey, and are better at stopping prey (e.g. Prokop & Grygláková 2005; Harmer *et al.* 2015). However, web investment was not constant across urbanisation gradients: at the landscape scale, community-averaged Capture Thread Length values increased with urbanisation, an effect driven by ITV (Figs 1, 2). There was no such effect at the local scale when ITV was accounted for (Figs 1, 2). The ITV-driven increase in web surface with landscape-level urbanisation, despite a decrease in mesh width, is likely linked to the ITV-driven increase in CTL along the same gradient. Similarly, the absence of overall changes in CTL with local urbanisation, in combination with the decrease in mesh width, likely explains the decrease of web surface with urbanisation at this spatial scale.

Overall, observed shifts in community web traits in response to urbanisation appear to be adaptive responses to prey spectrum, particularly prey size, changes. In particular, the ITV-driven increase in silk investment with landscape-level urbanisation allowed for simultaneous adaptive shifts in mesh width and web surface at this spatial scale, despite an existing trade-off between these two traits. Effects of species replacement and ITV mostly went in the same direction, but ITV generally accentuated responses (Fig. 2). The fact that spiders were less abundant in locally urbanised communities may indicate that these changes, however, do not fully compensate the strong decrease in prey availability, at least at this spatial scale.

Intraspecific variation in web characteristics has important consequences for prey control by spider communities in cities. Prey biomass intercepted daily decreased with increased urbanisation at both spatial scales and whether ITV in webs was considered in estimates or not. However, this decrease was less important at the landscape scale when ITV was accounted for (Figs 4, 5). By contrast, intraspecific variation had no significant effect on local scale responses. These results also hold for *per capita* measures of prey interception (Supplementary Figure 3), meaning they are not only caused by variation in spider abundance (or the lack thereof). Intraspecific trait variation may therefore buffer ecosystem functioning against the effects of urbanisation. This effect is likely mediated by CTL, as it is only observed at the landscape scale. These results are in line with predictions that high intraspecific trait variation should contribute to ecosystem function stability in changing environments when the relationship between trait and function depends on environmental conditions (Wright, Ames & Mitchell 2016), as is the case with orb-webs.

Consequences of urbanisation were scale-dependent: spider communities were less able to compensate for the negative effects of urbanisation on abundance, traits or function at the more local scale. Local scale conservation actions (in e.g. urban parks or gardens) may therefore provide opportunities to maintain biodiversity and ecosystem functioning in larger urban ecosystems (Philpott *et al.* 2013). This study highlights the importance of intraspecific variation in community responses to urbanisation and other forms of anthropogenic changes. Although trait shifts mediated by species turnover are observed, ignoring ITV can potentially lead to underestimates of community trait responses to environmental changes. In addition, the proportion of among-communities variation explained by ITV is not always a good indication of its ecological importance, as ITV can contribute disproportionately to responses to environmental gradients. By facilitating trait-environment matching, intraspecific trait variation is also an important component of ecosystem function resilience in the face of urbanisation. While we do acknowledge this may in some cases be too costly or time-consuming (but see Lepš *et al.* 2011), we therefore plead for a more systematic assessment of ITV contributions during trait-based analyses, especially in the case of behavioural traits or responses to human-induced environmental changes.

## Authors’ Contributions

DB and MD conceived the ideas and designed methodology (DB: general and sampling methodology; MD: analysis methodology); DB, JD and MDC collected the data; JD, MDC and MD analysed the data; MD led the writing of the manuscript. All authors contributed critically to the drafts and gave final approval for publication.

## Acknowledgements

We are grateful to Pieter Vantieghiem for his assistance during field sampling. Financial support came from the Belspo IAP project P7/04 SPEEDY (SPatial and environmental determinants of Eco-Evolutionary DYnamics: anthropogenic environments as a model). MD is a postdoctoral fellow funded by a grant from the Fyssen Foundation.

## Data Accessibility

Data will be made available on Dryad upon final acceptance

## References

Albert, C.H., Grassein, F., Schurr, F.M., Vieilledent, G. & Violle, C. (2011) When and how should intraspecific variability be considered in trait-based plant ecology? Perspectives in Plant Ecology, Evolution and Systematics, 13, 217–225.

Aronson, M.F.J., Sorte, F.A.L., Nilon, C.H., Katti, M., Goddard, M.A., Lepczyk, C.A., Warren, P.S., Williams, N.S.G., Cilliers, S., Clarkson, B., Dobbs, C., Dolan, R., Hedblom, M., Klotz, S., Kooijmans, J.L., Kühn, I., MacGregor-Fors, I., McDonnell, M., Mörtberg, U., Pyšek, P., Siebert, S., Sushinsky, J., Werner, P. & Winter, M. (2014) A global analysis of the impacts of urbanization on bird and plant diversity reveals key anthropogenic drivers. Proceedings of the Royal Society of London B: Biological Sciences, 281, 20133330.

Bates, D., Mächler, M., Bolker, B. & Walker, S. (2015) Fitting linear mixed-effects models using lme4. Journal of Statistical Software, 67.

Benjamini, Y. & Hochberg, Y. (1995) Controlling the false discovery rate: a practical and powerful approach to multiple testing. Journal of the Royal Statistical Society. Series B (Methodological), 57, 289–300.

Blackledge, T.A. & Eliason, C.M. (2007) Functionally independent components of prey capture are architecturally constrained in spider orb webs. Biology Letters, 3, 456–458.

Blackledge, T.A. & Zevenbergen, J.M. (2006) Mesh width influences prey retention in spider orb webs. Ethology, 112, 1194–1201.

Blamires, S.J. (2010) Plasticity in extended phenotypes: orb web architectural responses to variations in prey parameters. Journal of Experimental Biology, 213, 3207–3212.

Blamires, S.J., Thompson, M.B. & Hochuli, D.F. (2007) Habitat selection and web plasticity by the orb spider *Argiope keyserlingi* (Argiopidae): Do they compromise foraging success for predator avoidance? Austral Ecology, 32, 551–563.

Bonte, D., Lanckacker, K., Wiersma, E. & Lens, L. (2008) Web building flexibility of an orb-web spider in a heterogeneous agricultural landscape. Ecography, 31, 646–653.

Cornwell, W.K. & Ackerly, D.D. (2009) Community assembly and shifts in plant trait distributions across an environmental gradient in coastal California. Ecological Monographs, 79, 109–126.

Croci, S., Butet, A. & Clergeau, P. (2008) Does urbanization filter birds on the basis of their biological traits? The Condor, 110, 223–240.

Dray, S., Choler, P., Dolédec, S., Peres-Neto, P.R., Thuiller, W., Pavoine, S. & ter Braak, C.J.F. (2014) Combining the fourth-corner and the RLQ methods for assessing trait responses to environmental variation. Ecology, 95, 14–21.

Duckworth, R.A. (2008) The role of behavior in evolution: a search for mechanism. Evolutionary Ecology, 23, 513–531.

Eberhard, W.G. (2013) The rare large prey hypothesis for orb web evolution: a critique. Journal of Arachnology, 41, 76–80.

Entling, W., Schmidt-Entling, M.H., Bacher, S., Brandl, R. & Nentwig, W. (2010) Body size– climate relationships of European spiders. Journal of Biogeography, 37, 477–485.

Evans, S.C. (2013) Stochastic modeling of orb-web capture mechanics supports the importance of rare large prey for spider foraging success and suggests how webs sample available biomass. MSc dissertation, University of Akron.

Fischer, J.D., Cleeton, S.H., Lyons, T.P. & Miller, J.R. (2012) Urbanization and the predation paradox: the role of trophic dynamics in structuring vertebrate communities. BioScience, 62, 809–818.

Foelix, R. (2010) Biology of Spiders, 3rd edition. Oxford University Press, Oxford, UK.

Gregorič, M., Kuntner, M. & Blackledge, T.A. (2015) Does body size predict foraging effort? Patterns of material investment in spider orb webs. Journal of Zoology, 296, 67–78.

Harmer, A.M.T., Clausen, P.D., Wroe, S. & Madin, J.S. (2015) Large orb-webs adapted to maximise total biomass not rare, large prey. Scientific Reports, 5.

Heiling, A.M. & Herberstein, M.E. (1998) The web of Nuctenea sclopetaria (Araneae, Araneidae): relationship between body size and web design. The Journal of Arachnology, 26, 91–96.

Herberstein, M.E. (1997) The effect of habitat structure on web height preference in three sympatric web-building spiders (Araneae, Linyphiidae). The Journal of Arachnology, 25, 93–96.

Herberstein, M.E. & Tso, I.-M. (2000) Evaluation of formulae to estimate the capture area and mesh height of orb webs (Araneoidea, Araneae). Journal of Arachnology, 28, 180–184.

Hlivko, J.T. & Rypstra, A.L. (2003) Spiders reduce herbivory: nonlethal effects of spiders on the consumption of soybean leaves by beetle pests. Annals of the Entomological Society of America, 96, 914–919.

Jung, V., Albert, C.H., Violle, C., Kunstler, G., Loucougaray, G. & Spiegelberger, T. (2014) Intraspecific trait variability mediates the response of subalpine grassland communities to extreme drought events. Journal of Ecology, 102, 45–53.

Kaiser, A., Merckx, T. & Van Dyck, H. (2016) The Urban Heat Island and its spatial scale dependent impact on survival and development in butterflies of different thermal sensitivity. Ecology and Evolution, 6, 4129–4140.

Knop, E. (2016) Biotic homogenization of three insect groups due to urbanization. Global Change Biology, 22, 228–236.

Lajoie, G. & Vellend, M. (2015) Understanding context dependence in the contribution of intraspecific variation to community trait–environment matching. Ecology, 96, 2912–2922.

Lavorel, S. & Garnier, E. (2002) Predicting changes in community composition and ecosystem functioning from plant traits: revisiting the Holy Grail. Functional Ecology, 16, 545–556.

Lavorel, S., Grigulis, K., Lamarque, P., Colace, M.-P., Garden, D., Girel, J., Pellet, G. & Douzet, R. (2011) Using plant functional traits to understand the landscape distribution of multiple ecosystem services. Journal of Ecology, 99, 135–147.

Lepš, J., de Bello, F., Šmilauer, P. & Doležal, J. (2011) Community trait response to environment: disentangling species turnover vs intraspecific trait variability effects. Ecography, 34, 856–863.

Liu, Z., He, C., Zhou, Y. & Wu, J. (2014) How much of the world’s land has been urbanized, really? A hierarchical framework for avoiding confusion. Landscape Ecology, 29, 763–771.

Lowe, E.C., Wilder, S.M. & Hochuli, D.F. (2014) Urbanisation at multiple scales is associated with larger size and higher fecundity of an orb-weaving spider. PLoS ONE, 9, e105480.

Lubin, Y., Ellner, S. & Kotzman, M. (1993) Web relocation and habitat selection in desert widow spider. Ecology, 74, 1916–1928.

Ludy, C. (2007) Prey selection of orb-web spiders (Araneidae) on field margins. Agriculture, Ecosystems & Environment, 119, 368–372.

Mayntz, D., Toft, S. & Vollrath, F. (2003) Effects of prey quality and availability on the life history of a trap-building predator. Oikos, 101, 631–638.

McKinney, M.L. (2006) Urbanization as a major cause of biotic homogenization. Biological Conservation, 127, 247–260.

McKinney, M.L. (2008) Effects of urbanization on species richness: A review of plants and animals. Urban Ecosystems, 11, 161–176.

Miyashita, T., Shinkai, A. & Chida, T. (1998) The effects of forest fragmentation on web spider communities in urban areas. Biological Conservation, 86, 357–364.

Modlmeier, A.P., Keiser, C.N., Wright, C.M., Lichtenstein, J.L. & Pruitt, J.N. (2015) Integrating animal personality into insect population and community ecology. Current Opinion in Insect Science, 9, 77–85.

Moretti, M., De Bello, F., Roberts, S.P.M. & Potts, S.G. (2009) Taxonomical vs. functional responses of bee communities to fire in two contrasting climatic regions. Journal of Animal Ecology, 78, 98–108.

Oliveira, M.O., Freitas, B.M., Scheper, J. & Kleijn, D. (2016) Size and sex-dependent shrinkage of Dutch bees during one-and-a-half centuries of land-use change. PLOS ONE, 11, e0148983.

Opell, B.D., Berger, A.M. & Shaffer, R.S. (2007) The body size of the New Zealand orb-weaving spider Waitkera waitakerensis (Uloboridae) is directly related to temperature and affects fecundity. Invertebrate Biology, 126, 183–190.

Philpott, S.M., Cotton, J., Bichier, P., Friedrich, R.L., Moorhead, L.C., Uno, S. & Valdez, M. (2013) Local and landscape drivers of arthropod abundance, richness, and trophic composition in urban habitats. Urban Ecosystems, 17, 513–532.

Pickett, S.T.A., Cadenasso, M.L., Grove, J.M., Nilon, C.H., Pouyat, R.V., Zipperer, W.C. & Costanza, R. (2001) Urban ecological systems: linking terrestrial ecological, physical, and socioeconomic components of metropolitan areas. Annual Review of Ecology and Systematics, 32, 127–157.

Pigliucci, M. (2001) Phenotypic plasticity: beyond nature and nurture. Johns Hopkins University Press.

Prokop, P. & Grygláková, D. (2005) Factors affecting the foraging success of the wasp-like spider Argiope bruennichi (Araneae): role of web design. Biologia, 60, 165–169.

R Core Team. (2016) R: a language and environment for statistical computing. R Foundation for Statistical Computing, Vienna, Austria.

Renauld, M., Hutchinson, A., Loeb, G., Poveda, K. & Connelly, H. (2016) Landscape simplification constrains adult size in a native ground-nesting bee. PLOS ONE, 11, e0150946.

Riechert, S.E. & Maupin, J. (1998) Spider effects on prey: tests for superfluous killing in five web-builders. Proceedings of the 17th European Colloquium of Arachnology, pp. 203–210.

Roberts, M. (1993) The spiders of Great Britain and Ireland. Apollo Books, Stenstrup, Denmark.

Rypstra, A.L. & Buddle, C.M. (2013) Spider silk reduces insect herbivory. Biology Letters, 9, 20120948.

Sabo, J., Bastow, J. & Power, M. (2002) Length–mass relationships for adult aquatic and terrestrial invertebrates in a California watershed. Journal of the North American Benthological Society, 21, 336–343.

Sandoval, C.P. (1994) Plasticity in Web Design in the Spider Parawixia bistriata: A Response to Variable Prey Type. Functional Ecology, 8, 701–707.

Sattler, T., Borcard, D., Arlettaz, R., Bontadina, F., Legendre, P., Obrist, M.K. & Moretti, M. (2010) Spider, bee, and bird communities in cities are shaped by environmental control and high stochasticity. Ecology, 91, 3343–3353.

Scharf, I., Lubin, Y. & Ovadia, O. (2011) Foraging decisions and behavioural flexibility in trap-building predators: a review. Biological Reviews, 86, 626–639.

Schneider, J.M. & Vollrath, F. (1998) The effect of prey type on the geometry of the capture web of Araneus diadematus. Naturwissenschaften, 85, 391–394.

Sensenig, A., Agnarsson, I. & Blackledge, T.A. (2010) Behavioural and biomaterial coevolution in spider orb webs. Journal of Evolutionary Biology, 23, 1839–1856.

Seto, K.C., Güneralp, B. & Hutyra, L.R. (2012) Global forecasts of urban expansion to 2030 and direct impacts on biodiversity and carbon pools. Proceedings of the National Academy of Sciences, 109, 16083–16088.

Sherman, P.M. (1994) The orb-web: an energetic and behavioural estimator of a spider’s dynamic foraging and reproductive strategies. Animal Behaviour, 48, 19–34.

Siefert, A., Violle, C., Chalmandrier, L., Albert, C.H., Taudiere, A., Fajardo, A., Aarssen, L.W., Baraloto, C., Carlucci, M.B., Cianciaruso, M.V., de L. Dantas, V., de Bello, F., Duarte, L.D.S., Fonseca, C.R., Freschet, G.T., Gaucherand, S., Gross, N., Hikosaka, K., Jackson, B., Jung, V., Kamiyama, C., Katabuchi, M., Kembel, S.W., Kichenin, E., Kraft, N.J.B., Lagerström, A., Bagousse-Pinguet, Y.L., Li, Y., Mason, N., Messier, J., Nakashizuka, T., Overton, J.M., Peltzer, D.A., Pérez-Ramos, I.M., Pillar, V.D., Prentice, H.C., Richardson, S., Sasaki, T., Schamp, B.S., Schöb, C., Shipley, B., Sundqvist, M., Sykes, M.T., Vandewalle, M. & Wardle, D.A. (2015) A global meta-analysis of the relative extent of intraspecific trait variation in plant communities. Ecology Letters, 18, 1406–1419.

Sih, A., Stamps, J., Yang, L.H., McElreath, R. & Ramenofsky, M. (2010) Behavior as a key component of integrative biology in a human-altered world. Integrative and Comparative Biology, 50, 934–944.

Simons, N.K., Weisser, W.W. & Gossner, M.M. (2016) Multi-taxa approach shows consistent shifts in arthropod functional traits along grassland land-use intensity gradient. Ecology, 97, 754–764.

Smith, R.M., Gaston, K.J., Warren, P.H. & Thompson, K. Urban domestic gardens (VIII) ⃞: environmental correlates of invertebrate abundance. Biodiversity & Conservation, 15, 2515–2545.

United Nations Population Division. (2015) World urbanization prospects: the 2014 revision. United Nations, Department of Economic and Social Affairs, New York, USA.

Venner, S., Thevenard, L., Pasquet, A. & Leborgne, R. (2001) Estimation of the web’s capture thread length in orb-weaving spiders: determining the most efficient formula. Annals of the Entomological Society of America, 94, 490–496.

Violle, C., Enquist, B.J., McGill, B.J., Jiang, L., Albert, C.H., Hulshof, C., Jung, V. & Messier, J. (2012) The return of the variance: intraspecific variability in community ecology. Trends in Ecology & Evolution, 27, 244–252.

Vriens, L., Bosch, H., De Knijf, S., De Saeger, S., Guelinckx, R., Oosterlynck, P., Van Hove, M. & Paelinckx, D. (2011) De Biologische Waarderingskaart - Biotopen En Hun Verspreiding in Vlaanderen En Het Brussels Hoofdstedelijk Gewest [En: The Biological Valuation Map - Habitats and Their Distribution in Flanders and the Brussels Capital Region]. Mededelingen van het Instituut voor Natuuren Bosonderzoek, Brussels, Belgium.

Wong, B.B.M. & Candolin, U. (2015) Behavioral responses to changing environments. Behavioral Ecology, 26, 665–673.

Wright, J.P., Ames, G.M. & Mitchell, R.M. (2016) The more things change, the more they stay the same? When is trait variability important for stability of ecosystem function in a changing environment. Philosophical Transactions of the Royal Society of London B: Biological Sciences, 371, 20150272.

Zschokke, S. (1999) Nomenclature of the Orb-Web. The Journal of Arachnology, 27, 542–546.

